# Comparison of Proton Transfer Paths to the Q_A_ and Q_B_ Sites of the *Rb. sphaeroides* Photosynthetic Reaction Centers

**DOI:** 10.1101/2022.02.10.479886

**Authors:** Rongmei Judy Wei, Yingying Zhang, Junjun Mao, Divya Kaur, Umesh Khaniya, M.R. Gunner

## Abstract

The photosynthetic bacterial reaction centers from purple non-sulfur bacteria use light energy to drive the transfer of electrons from cytochrome c to ubiquinone. Ubiquinone bound in the Q_A_ site cycles between quinone, Q_A_, and anionic semiquinone, Q_A_^•-^, being reduced once and never binding protons. In the Q_B_ site, ubiquinone is reduced twice by Q_A_^•-^, binds two protons and is released into the membrane as the quinol, QH_2_. The network of hydrogen bonds formed in a molecular dynamics trajectory was drawn to investigate proton transfer pathways from the cytoplasm to each quinone binding site. Q_A_ is isolated with no path for protons to enter from the surface. In contrast, there is a complex and tangled network requiring residues and waters that can bring protons to Q_B_. There are three entries from clusters of surface residues centered around HisH126, GluH224, and HisH68. The network is in good agreement with earlier studies, Mutation of key nodes in the network, such as SerL223, were previously shown to slow proton delivery. Mutational studies had also shown that double mutations of residues such as Asp M17 and AspL210 along multiple paths in the network presented here slow the reaction, while single mutations do not. Likewise, mutation of both HisH126 and HisH128, which are at the entry to two paths reduce the rate of proton uptake.

## 1. Introduction

Intra-protein reactions that gain or lose protons require proton transfer to the active site. The membranes of mitochondria, chloroplasts and bacteria contain many proteins that carry out a series of electron transfer reactions that remove protons from one side of the membrane and release them to the other. Membranes thus have a more negative, N-side that is the source of protons and a positive, P-side where protons are released. The resulting transmembrane electrochemical gradient fuels production of ATP (Mitchell 1976; Mulkidjanian 2007; Walker 2013) and other necessary cellular functions.

In oxygenic photosynthesis, the electron transfer chain moves electrons from water to NADH to make metabolically useful reduced products such as sugar. PSII (Kern and Renger 2007; Shen 2015; Vinyard and Brudvig 2017) and the b_6_f complex (Darrouzet et al. 2004) add to the proton gradient by placing their oxidation reactions, which release protons, on the lumen, P-side of the membrane (Umena et al. 2011; Cox et al. 2013; Vinyard and Brudvig 2017; Pantazis 2018). Their reduction reactions, that are coupled to proton binding, are on the stromal, N-side of the membrane. Thus, the lumen is acidified and stroma made more basic without protons flowing through the protein (Cardona et al. 2012; Müh et al. 2012).

In the aerobic mitochondrial or bacterial electron transfer chain, electrons from reduced metabolites such as NADH and succinate pass through a series of proteins to reduce O_2_ in cytochrome c oxidase. Protons are pumped through Complex I (Baradaran et al. 2013) and cytochrome c oxidase (Kaila et al. 2010) from N- to P-side through well-studied, gated proton transfer paths (Lee et al. 2012; Kaila 2018). The bc_1_ complex (Rich 2004; Cape et al. 2006; Mulkidjanian 2010; Sarewicz and Osyczka 2015), in the same protein family as the chloroplast b_6_f complex, adds to the proton gradient by separating the oxidation and reduction chemistry to the more positive and negative sides of the membrane.

The anoxygenic photosynthetic bacteria, that are our subject here, add to the proton gradient by coupling the bacterial reaction centers (RCs) and cytochrome bc_1_ complex in a light driven electron transfer cycle. The overall reaction in RCs is

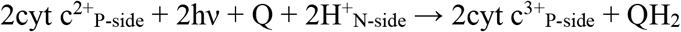

The quinol product then reduces the oxidized cytochrome c in the bc_1_ complex to add to the proton gradient. Thus, the only product of the combined action of RCs and the bc_1_ complex is to carry out light driven addition to the proton gradient (Lin et al. 1994; Venturoli et al. 1998; Okamura et al. 2000c; Mulkidjanian 2007; Lancaster et al. 2007).

### Quinone reactions in bacterial RCs

Excitation of a dimer of bacteriochlorophyll, P, initiates electron transfer across the protein, via the L branch bacteriopheophytin to Q_A_ forming the semiquinone Q_A_^•-^, which then reduces Q_B_ (Figure 1, eqn. 1a). P^+^ is reduced back to the ground state by cytochrome c.

**Figure 1:**
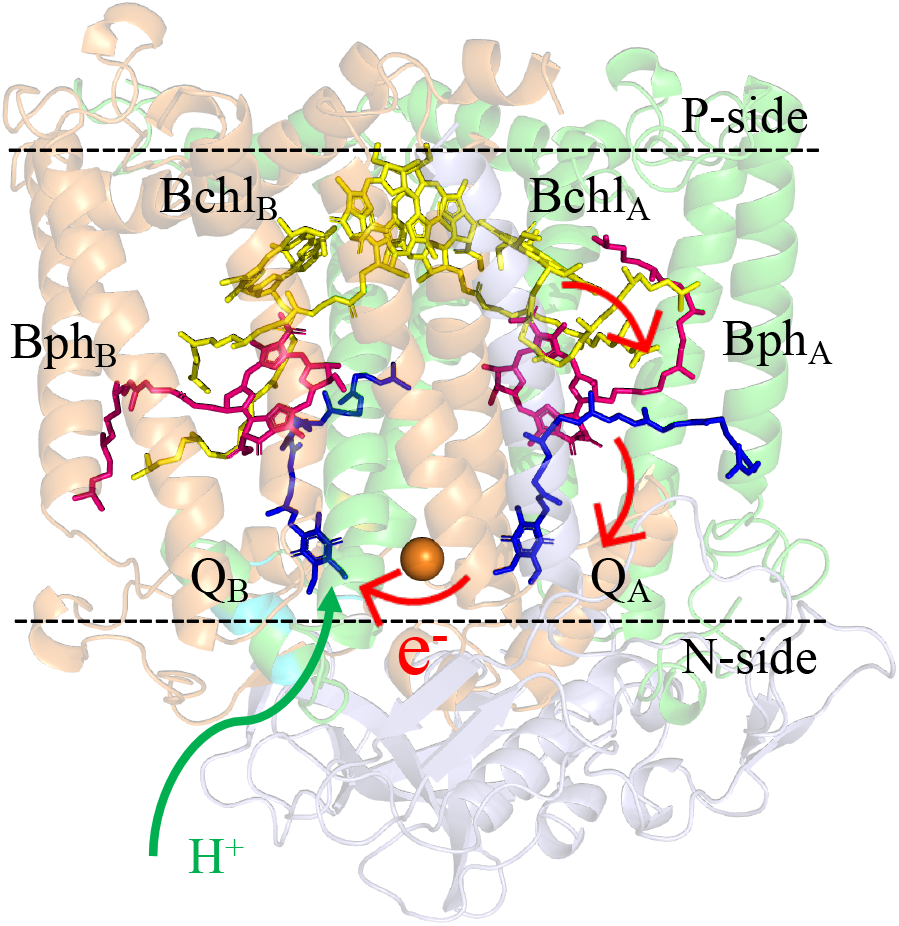
The purple non-sulfur bacterial photosynthetic reaction centers of *Rb. sphaeroides*, PDB ID: 1AIG (Stowell et al. 1997). L, M, and H subunits are in green, orange and purple, respectively. The dashed lines show the approximate location of the membrane surface. Red arrows indicate the electron transfer route, and the green arrow shows where protons move from the N-side, cytoplasm to Q_B_. Quinones are in blue, bacterialpheophytins (BPh) are in magenta, and bacterialchlorophylls (BChl) are in yellow.

The second excitation leads to formation of Q_A_^•-^Q_B_^•-^ (Eqn 1b). Uphill proton binding by the Q_B_^•-^ occurs before electron transfer from Q_A_^•-^. Thus, Q_A_^•-^ reduces the neutral semiquinone Q_B_H^•^ to the anionic quinol Q_B_H^-^. A second proton is bound forming Q_B_H_2_ which is released into the membrane (Graige et al. 1996).

Sequence of electron and proton transfer reactions at the acceptor side of RCs:

First reduction:

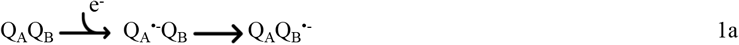

Second reduction:

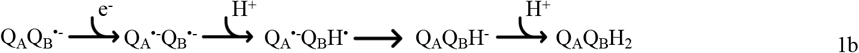

Thus, two protons are bound to Q_B_, one to each carbonyl, removing protons from the cytoplasm. The quinol is released to the quinone pool and a new quinone will be bound to the Q_B_ site. In contrast, Q_A_ never leaves the protein. This same sequence of reactions is carried out by the quinones on the stromal side of PSII (Robinson and Crofts 1983; Müh et al. 2012; Saito et al. 2013). The P-side surface of bacterial RCs is capped by the H-subunit, which is missing in PSII (Okamura et al. 2000c; Sakashita et al. 2017). Thus, in RCs Q_A_ and Q_B_ are deeply buried in the protein requiring long-range proton transfer to Q_B_, while the quinones in PSII are much closer to the surface.

### Coupling of proton transfers and Q_B_ reduction

The pK_a_ of Q_B_^•-^ is low enough that no proton is bound directly to the semiquinone (Graige et al. 1999; Gunner et al. 2008; Maróti et al. 2015). However, protons are bound into the protein to help stabilize the anionic semiquinone and to bring a proton to a position where it can be transferred to Q_B_^•-^ prior to the second reduction (McPherson et al. 1988; Maróti and Wraight 1997; Alexov and Gunner 1999). Under standard conditions protonation of Q_B_^•-^ is always faster than electron transfer so proton transfer is never rate limiting (Okamura et al. 2000c; Alexov et al. 2000; Wraight 2004).

### Requirements for proton transfer through proteins

Soluble or membrane proteins, requiring protons at buried reaction sites need proton transfer paths. Proton transfer (PT) differs from the transfer of other substrates in that the proton that enters the protein is not the one that arrives at the active site. Protons exchange, as they are passed through networks of water molecules and polar, acidic and basic residues. The transported proton is never free but is associated with water as hydronium, or larger complexes (Wraight 2006; Farahvash and Stuchebrukhov 2018), or bound to acidic or basic residues.

Protons are passed via the Grotthuss mechanism where each group in a hydrogen bonded chain transiently gains a H^+^ from an upstream donor and loses one to the downstream acceptor (Swanson et al. 2007; Agmon et al. 2016). Thus, each H^+^ shifts ownership by one group, while a proton enters one side of the chain and another exits the other end. Grotthuss PT requires a preorganized hydrogen bond chain and following the exit of a proton, the hydrogen bonded chain must rearrange to let the next one go through (Agmon 1995; Cukierman 2006; DeCoursey and Hosler 2014). Grotthuss competent residues, such as water and hydroxyl groups (Ser, Thr, Tyr), neutral His, protonated carboxylic acids, and neutral Lys are both a hydrogen bond acceptor and donor. Ionized acidic and basic residues Asp^-^, Glu^-^. His^+^ and Lys^+^ can be proton loading sites (PLS). PLS facilitate proton transfer by transiently binding or loosing protons as reaction intermediates. Residues we denote as fences are unlikely to serve in Grotthuss PT or as PLS, include Asn, Gln, Trp and Arg. Arg is considered a fence residue as its proton affinity is too high to easily change protonation in a PLS (Fitch et al. 2015) and its geometry requires a proton lost from one nitrogen to be replaced to the same one, so it cannot help protons move forward easily (Ge and Gunner 2016).

### Structures of proton transfer paths

Proton transfer paths are identified by finding hydrogen bonded networks of water molecules and Grotthuss competent and acidic and basic amino acids. Proton transfer paths through RCs (Okamura and Feher 1992; Okamura et al. 2000c), PSII (Bondar and Dau 2012; Kaur et al. 2021), Cytochrome c oxidase (Cai et al. 2018), Complex I (Khaniya et al. 2020) and other proteins (Siemers et al. 2019; Bertalan et al. 2020) have been investigated. Proton transfer networks differ in the participation of different types of residues. Some long-range PT paths have many intervening residues (Cai et al. 2018; Khaniya et al. 2020). In contrast, proton release from the Oxygen Evolving Complex in PSII to the lumen can travel ≈15Å via water filled channels without needing to hop on and off intervening amino acid side chains (Röpke et al. 2020; Kaur et al. 2021). Networks also differ in their structures. Linear proton transfer networks can be found where single residues stand at the entrance and exit with a well-define channel between them. Examples include the antiporter channels in complex I (Di Luca et al. 2017), the D and K channels in cytochrome c oxidase (Cai et al. 2020) and the broad, narrow and large channels for proton exit on the lumen side of PSII (Kaur et al. 2021). In contrast, complex pathways have multiple entries, exits and a tangled network of possible routes for proton transfer. Examples include, the P-side proton exit in cytochrome c oxidase (Cai et al. 2018), the E-channel of complex I (Khaniya et al. 2020) and the ≈10Å region around the OEC that connects the proton release sites to the three possible long-range exit channels (Kaur et al. 2021). While linear paths are often obvious in the protein structure, complex paths are difficult to visualize as they can extend through a large portion of the protein.

### Proton transfer paths in RCs

RCs from the purple photosynthetic bacteria *Rps. viridis* were the first membrane protein with an atomic resolution crystal structure (Deisenhofer et al. 1985). With this information and subsequent structures, the *Rb. sphaeroides* RCs served as a model protein for the study of photosynthesis. Many experiments (Paddock et al. 1989; Okamura et al. 2000c; Wraight 2004; Ptushenko et al. 2015) and simulations (Beroza et al. 1995; Rabenstein et al. 1998, 2000; Hammes-Schiffer 2001; Ishikita et al. 2003; Ishikita and Knapp 2004a, 2005) studies have helped us understand the rules for the proton uptake coupled to reduction of Q_A_ and Q_B_. These earlier studies recognized that there are many acidic and basic residues near Q_B_ that modulate the quinone reduction potential (Paddock et al. 1994a, 1995; Alexov et al. 2000), contribute to the storage of protons within the protein that can be passed to Q_B_- and to Q_B_H^-^ (Okamura and Feher 1992) and form the pathway for proton transfer. Mutations were made of many residues to determine their effect on the quinone electron and proton transfer reactions (Okamura et al. 2000c).

The work presented here will provide a new analysis of the possible proton transfer paths around both Q_A_ and Q_B_. All hydrogen bonds around the quinones are found in a molecular dynamics trajectory carried out on *Rb. sphaeroides* RCs. Network analysis identifies all possible paths. Three entries are found leading to Q_B_ via a tangled web of hydrogen bonded connections. The residues contributing to different paths are identified and compared with the influence of previous mutations of the protein. In contrast, even with a well-hydrated RC structure Q_A_ remains insulated from the solvent by its burial in a largely hydrophobic region.

## 2. Methods

### 2.1 MD simulations

The calculations start with the 2.6 Å resolution X-ray crystal structure of *Rhodobacter (Rb.) sphaeroides* RCs (PDB ID: 1AIG) with Q_B_ in the proximal binding site (Stowell et al. 1997). The L, H and M subunits and all cofactors are included. The system was prepared using the CHARMM-GUI bilayer membrane builder, embedding the protein in a bilayer of 1-palmitoyl-2-oleoyl-sn-glycero-3-phosphocholine (POPC) (Jo et al. 2008). The CHARMM36 force field is used for the protein and lipids (Klauda et al. 2010) and TIP3P parameters for water molecules (Jorgensen et al. 1983; Neria et al. 1996; Lee et al. 2016). The CHARMM force field parameters for POPC and RC cofactors ubiquinone, bacteriochlorophyll, bacteriopheophytin and non-heme, Fe^2+^ are from Matyushov (Ceccarelli et al. 2003; LeBard et al. 2008). The parameters of cofactors are patched to the protein. HisL190, HisL230, HisM219, HisM266 and GluM234 are connected to the non-heme iron and a His (HisL153, HisM202, HisL173 and HisM182) to each of the four bacteriochlorophylls. With thanks to Matyuskov the RC cofactor parameters are available at https://github.com/GunnerLab/mcce-toolbox/tree/master/hb_analysis_MD. The RC is assembled in a 115 Å × 115 Å × 115 Å cubic box with ≈30,000 TIP3P water molecules and ≈300 lipid molecules in the membrane. 96 Na^+^ and 84 Cl^-^_are added to maintain charge neutrality at 150 mM ionic strength. There are 188,508 atoms (Figure SI.1A).

#### Assigning residue protonation states

The protonation states of all residues are determined with MCCE (Song et al. 2009) using methods and parameter described previously (Kaur et al. 2021). MCCE allows each acidic and basic residue to choose between protonated and deprotonated states via Monte Carlo sampling. One microstate of a protein is a snapshot with all residues with one assigned protonation state. The most probable microstate is used as input to fix the RC protonation state in MD. MCCE supports the CHARMM default protonation states with Asp and Glu deprotonated, Arg and Lys protonated and His are neutral with the proton on ND1 (HSD) for most residues. However, AspL210, GluH43, GluL104, GluM236 are protonated so neutral; while HisH126, HisH128 and HisM301 are protonated so positively charged. An additional 2.81 Å distance restraint was enforced with an energy of 1000 Kcal/mol/Å to retain the Q_B_ Oxygen to HisL190 Nitrogen hydrogen bond. In the absence of the constraint, Q_B_ will move into the proximal position (Stowell et al. 1997) (Figure SI.2).

### 2.2. OpenMM MD simulation

The MD simulations were done using OpenMM (Eastman et al. 2017) and are performed with periodic boundary conditions at constant pressure (1 bar) and temperature (303.15 K) using Langevin dynamics with Nose-Hoover Langevin piston. The Particle Mesh Ewald (PME) algorithm is used for computing long-range electrostatic interactions. To relax the initial clashes, the system is equilibrated using OpenMM where the protein backbone, side chains and lipids are allowed to relax for 6450 ps. A time step of 2 fs is used. The production run is 150 ns long with snapshots saved every 10 ps. The snapshots from the 51-100 ns period are chosen for analysis as this has a stable backbone RMSD Figure SI.1B.

#### Finding the hydrogen bond network

The hydrogen bond network is mapped to follow the protons transfer path from solvent to Q_B_ and to compare the networks around Q_A_ and Q_B_. MDTraj (McGibbon et al. 2015) extracts each snapshot, providing 50 snapshot/ns. For each frame, the water molecules outside the protein are stripped off until only those with surface accessibility < 20% remain.

#### Finding hydrogen bonds

All hydrogen bonds between waters and Ser, Thr and Tyr, Asp, Glu, His and Lys side chains are found. These residues can participate in Grotthuss proton transfer or serve as proton loading sites. Hydrogen bonds have a distance of 1.2 Å to 3.5 Å between the donor and acceptor and the angle between donor hydrogen and acceptor is > 90°. The hydrogen bond network between residues via waters are found with the depth-first-search (DFS) algorithm. DFS traverses a graph structure from an arbitrary starting node, exploring as far as possible along each branch then backtracking along all paths (Mercado et al. 2021). In our hydrogen bond analysis tool, a random residue from the residue list is chosen as a starting node, then all the routes from this node that connect to Q_B_ are recorded. Backtracking routes enumerate all possible paths. This process is repeated until all residues have been subjected to DFS.

The trajectories are grouped into 1-50ns, 51-100ns, 101-150ns, denoted as 50, 100, 150, respectively. The network for each time block includes 2,500 frames. Residues are often connected via mobile water molecules, which are counted as maintaining the connection even if the waters have exchanged between frames. Cytoscape is used to visualize the network (Shannon et al. 2003). Networks are identified showing residue connections with no intervening waters (0w), a maximum or 1, 2, 3 or 4 intervening waters (denoted 0w, 1w, 2w, 3w, 4w). The exposure of each residue is calculated by MCCE tools that adds the atom solvent accessible surface area (SASA) for each atom in a residue. Surface exposed residues are defined as having more than 20% of their area touching solvent.

## 3. Results

### Residues in the hydrogen bond network leading to Q_B_

The hydrogen bond networks around Q_A_ and Q_B_ are derived using 50 ns time blocks from the MD trajectory carried out on *Rb. sphaeroides* RCs. The networks are built of waters and only residues that can participate in proton transport via the Grotthuss mechanism (Ser, Thr, Tyr and neutral His) or as proton loading sites (Asp, Glu, Lys and His). In drawing the network of residues, different numbers of intervening waters are included. Thus 100-0w shows only residues that are directly connected to one another in the block of the trajectory from 51-100 ns. Networks are then drawn using the same input atomic coordinates but now allowing residues to be connected by one intervening water (1w) or with 2, 3 and 4 waters (2w, 3w, and 4w).

Table 1 lists the amino acids in the hydrogen bond networks near Q_B_. For example, there are 26 residues in the 100-2w network, which analyzes the time block from 51-100 ns with a maximum of two intervening waters. These residues will be shown to provide multiple paths from the surface to Q_B_ via relatively few steps. The network is predominantly acidic with five Asp and Glu and three Lys. There are five His in the network. MCCE calculates that HisH126 and HisH128 are protonated and the other neutral in the most probable RC protonation microstate. GluM234 and His L190 are ligands to the non-heme iron bridging the two quinones so are unlikely to be available to transfer protons.

**Table 1:**
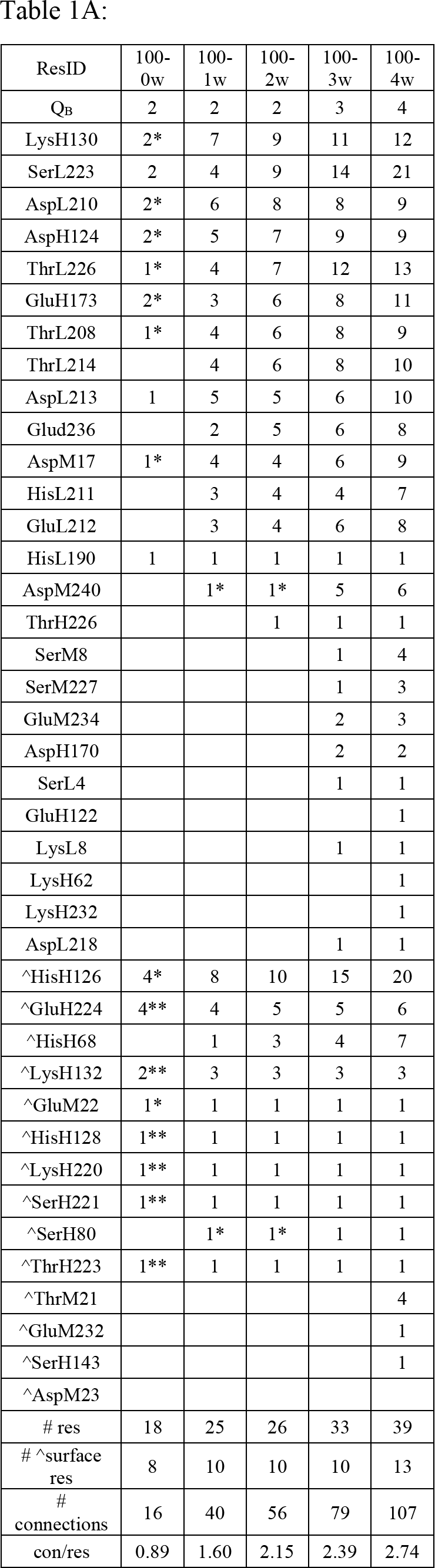
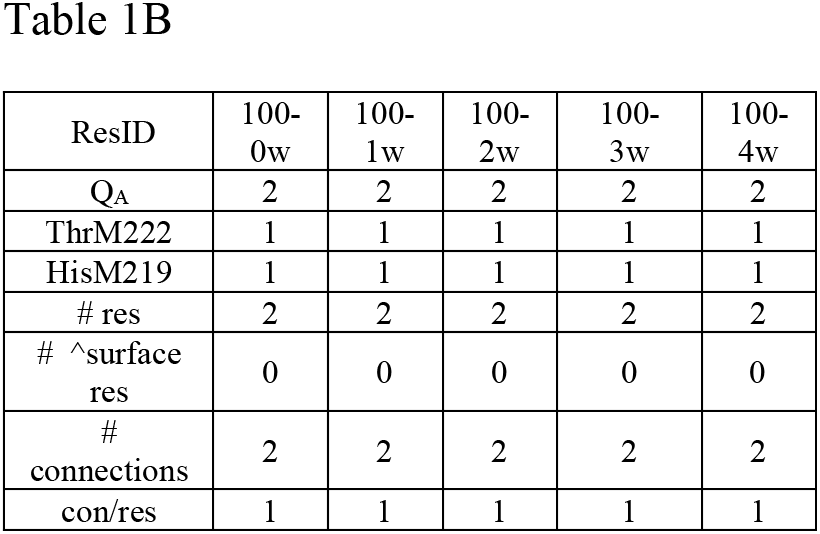
Count of connections between residues in the network connecting Q_B_ and Q_A_ to the surface with different number of intervening water molecules in the 51-100 ns time block of the trajectory. The number of other residues is a sum of connections in all of the 2,500 frames. Thus, each residue can make hydrogen bonds to different partners in different frames. The sums at the bottom of the table count the total number of connected residues in the network and the residues with 20% surface exposure. It provided the total number of connections in the network and the average number of connections per residue. ^ marks surface residues. Comparison of the 2w connections in the earlier and later time blocks are given in Table SI.3. Table 1A: Residues connections around Q_B_. With zero intervening waters considered (100-0w) two small networks are found that are not directly connected to Q_B_ (Fig 3a). They are marked as *: nearer Q_B_ and **: nearer the surface. Table 1B: Residues connections around Q_A_.

Table 1 shows how many residues each side chain that contribute to proton transfer is connected to in the network. Some residues are connected to more others than they have available hydrogen bond opportunities. This is because connections are noted when residues are hydrogen bonded in any of the 2,500 frames. Thus, residues can move to participate in multiple connections. For example, HisH126 can be connected via 0, 1 or 2 waters to 10 other resides including AspL210, AspM17, AspH124 and surface residue GluM22, while GluH224 can connect to ThrH223, SerH221, LysH220, and LysH132. In contrast, seven residues connect to only a single other because their positions are relatively fixed, they make hydrogen bonds in only a few frames or the waters needed for the connections are not always present. Overall, there are 56 connections made with an average of 2.15 connections/residue when two is the maximum number of intervening waters. Table SI.3 also shows the 2w network connections in the time blocks of the trajectory from 1 to 50 ns and from 101 to 150 ns. The network is well conserved. The highly interconnected resides remain so. A small number of residues, connected to only a single other move in and out of the network.

### The graphical network representation

A hydrogen bond network is drawn to show the connections between the residues. Figure 2 shows the residues in the 100-2w network in a physical and network representation. In the network the nodes are amino acids whose side chains are hydrogen bonded to another side chain via 0, 1 or 2 waters. The lines show the connections. Waters are shown as red dots in the physical representation, but not shown in the network.

**Figure 2:**
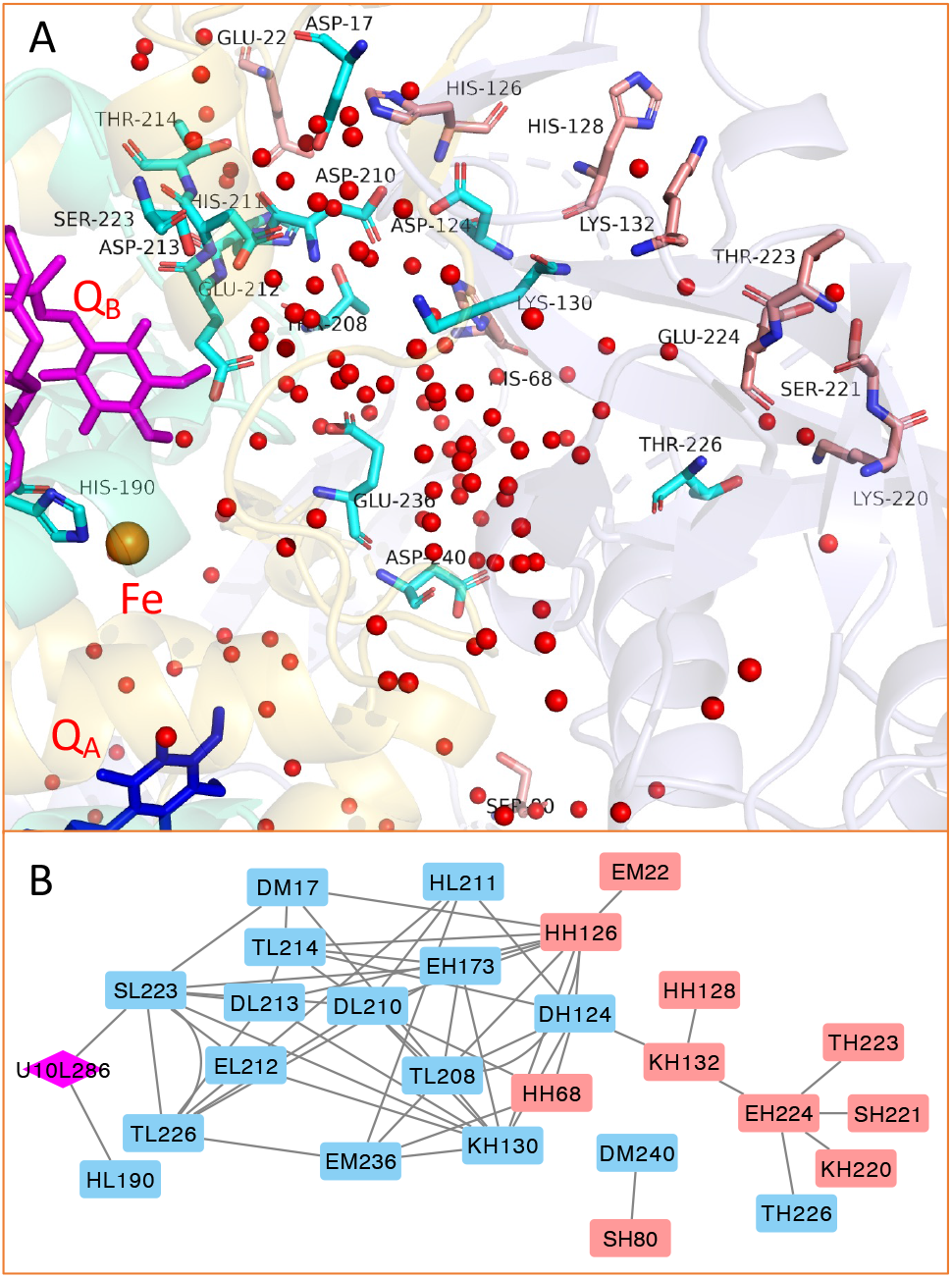
The hydrogen bond network obtained from the analysis of the trajectory from 51-100 ns showing connected by a maximum of two waters. Residues shown can facilitate proton transfer via a Grotthuss mechanism (Ser (S), Tyr (Y), Thr (T), neutral His (H)) or by transiently loading and unloading protons (Asp (D), Glu (E), Lys (K) and His (H)). Q_B_ is magenta; the surface residues are red-orange, and all other resides are blue. (A) Waters are red balls. Residues are in the L subunits with the exception of His126, His128, His68, Thr223, Glu224, Ser221, Lys220, Thr226, Lys132, Lys130, Glu173, Ser80 and Asp124, which are in the H subunit and Asp17, Glu236, Asp240 and Glu22, which are in the M subunit. (B) Nodes are residues denoted by one letter residue designation, chain (L, M, H) and residue ID. Lines are connections. The bridging waters are not shown.

### Comparison of networks with different numbers of waters bridging residues

Figure 3 shows the network drawn counting connections between different numbers of intervening waters. The 0w network shows connections only between residues that are directly hydrogen bonded to each other. Only HisL190 and SerL223 make connections to Q_B_. The network cannot go beyond AspL213 without the participation of water. The 0w network shows two additional small clusters that are not connected to Q_B_, but will be part of the extended network when connecting waters are considered.

**Figure 3.**
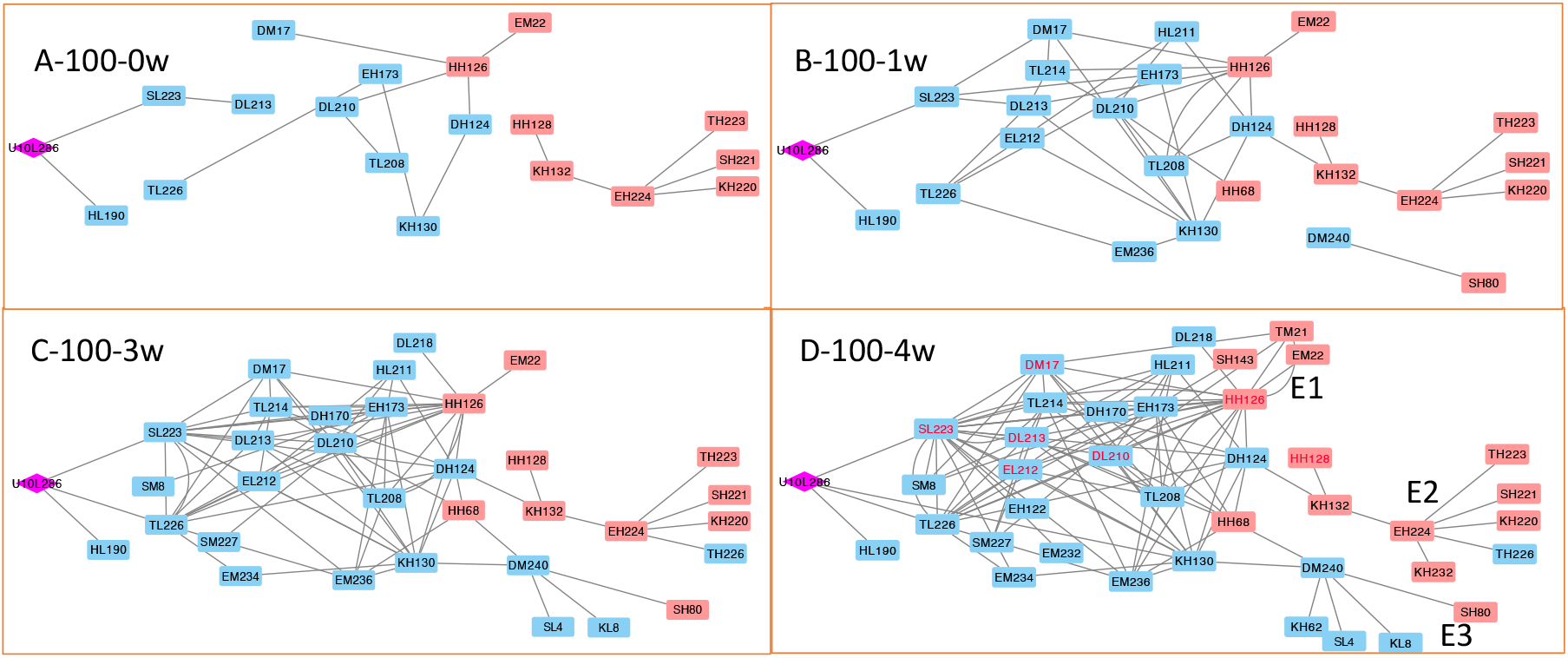
Network leading to Q_B_, The network is drawn using the same layout and colors as found in Fig 2B. All networks drawn using the 51-100ns block of the trajectory. (A) 0w requiring direct hydrogen bonding between residues; (B-D) allowing connections via (B) 1 water, (C) 3 waters or (D) 4 waters. Residues are retained on the network diagram if they are found in the 4w network and are connected to at least one other residue even if they have lost the connection to Q_B_. Thus, smaller, disconnected regions are shown in the 0w and 1w network.

Using the same input trajectory but counting connections between residues through one intervening water multiple paths from the surface to Q_B_ are seen. As we include more waters between residues the network becomes highly interconnected. There are 39 residues in the 4w network with an average of 2.74 connections between residues (Table 1).

### Entries to the network from the cytoplasm

Three separated clusters of surface exposed residues can lead into the Q_B_ site (Figure 3). In the 100-4w network, which is the most complete, Entry 1 is centered at HisH126 and connected to surface residues SerH143, ThrM21 and GluM22; Entry 2 is centered at GluH224, and connected toThrH223, SerH221, HisH128 and Lys H220, H232 and H132. Entry 3 has two surface residues SerH80 and HisH68 that are connected to each other via the buried AspM240. The shortest proton transfer paths from each entry go through three to four residues. For example, in the 51-100ns-4w network paths include:

Entry 1: HisH126→AspM17 → 1w → SerL223 → Q_B_.

Entry 2: LysH132 → 1w → AspH124 →3w → SerL223 → Q_B_.

Entry 3a: HisH68 →3w →AspL210 →2w →AspL213 → SerL223 → Q_B_.

Entry 3b: SerL80 → 3w →AspM240 →3w → LysH130→2w → SerL223 → Q_B_.

While many residues are involved in the network, all paths connect to Q_B_ via SerL223.

### The hydrogen bond network around Q_A_

Q_A_ and Q_B_ are symmetrically arranged around the non-heme iron. However, their functional role is quite different. Q_A_ is reduced only to the semiquinone and does not bind protons. In the hydrogen bond network, we find Q_A_ only connects to two side chains ThrM222 and the iron ligand HisM219. Remarkably the network does not expand when hops via waters are counted (Table 1B). Thus, within the 150 ns trajectory, no hydrogen bond network is found to connected Q_A_ to the surface (Figure 4).

**Figure 4.**
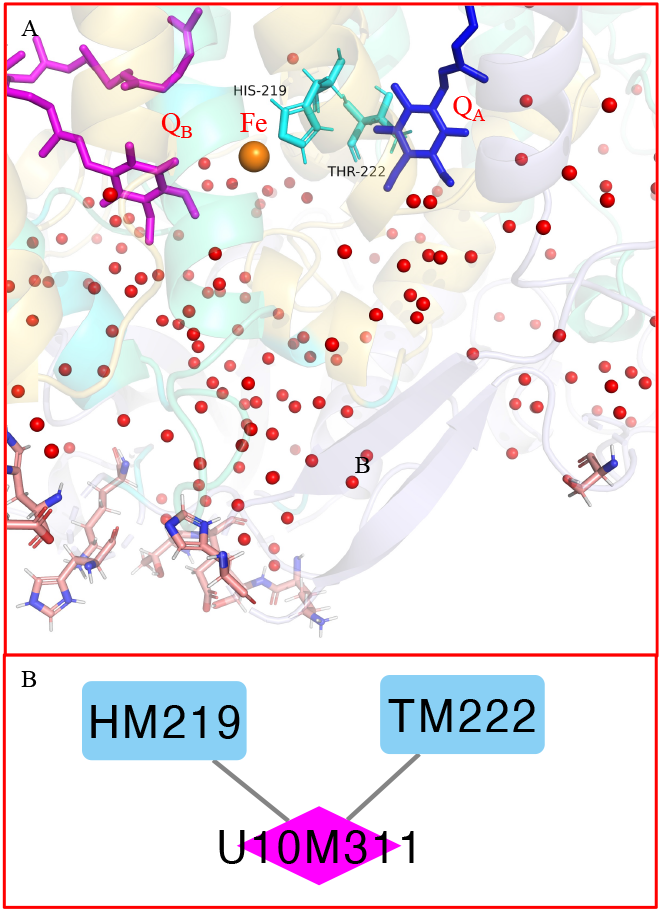
Hydrogen bond network connected to Q_A_. (A) Q_A_ is blue, Q_B_ is magenta, the surface residues are red-orange. Waters are red balls. (B) Nodes are residues denoted by residue name, chain name and residue ID. Lines are connections. The bridging waters are not shown. The network remains unchanged through the full 150 ns trajectory. Allowing connections through waters does not lead to additional connections to Q_A_.

## 4. Discussion

The hydrogen bond network around Q_A_ and Q_B_ were explored in an MD trajectory of *Rb. sphaeroides* RCs. Direct hydrogen bonds between residues form three small networks (Figure 3 Table 1). Q_B_ is directly connected to HisL190, a ligand to the non-heme iron, and to SerL223 which connects to AspL213. However, by adding transfers via only one bridging water the fundamental network from the surface to Q_B_ is established. As longer water paths are allowed the network becomes highly interconnected with many competing paths from the surface. Thus, proton transfers to Q_B_ appear to include active participation of both residues and waters. In contrast to Q_B_, Q_A_ is insulated from waters and other residues, with no path to the surface (Figure 4 Q_A_ network, Table 2).

Previous studies have been made of proton transfer paths to Q_B_ using inspection of the crystal structure (Okamura et al. 2000a; Paddock et al. 2003) and more detailed cluster based analysis (Krammer et al. 2009a, b). Paddock and Okamura (Abresch et al. 1998; Okamura et al. 2000a; Paddock et al. 2003) proposed three paths. P1 is a 20Å long pathway that connects Q_B_ via GluL212 to the surface at AspH124 and GluH224 or AspM240. P2 is also ≈20Å long, moving to Q_B_ via SerL223 from TyrM3 to GluH173. P3 is only ≈7 Å long with Q_B_ connecting to the surface residues HisH126 and HisH128 via AspM17 and AspL213.

The network presented here, obtained by an unbiased search of the hydrogen bonds formed in the MD trajectory, agree with earlier views of the proton transfer paths. The residues SerL223, AspL213, GluL212, AspM17, HisH126, and HisH128, whose importance had been shown earlier are key nodes in our network. However, there are some differences. For example, our the network finds three separate entries from the cytoplasm denoted E1, E2 and E3, but these are not fully aligned with the previously suggested entries to P1, P2, and P3. For example, HisH126 and HisH128 are in the entries E1 and E2 here, while they are both lead to P3 (Okamura et al. 2000a; Paddock et al. 2003). Additionally, the earlier work showed a connection in P3 between AspL213 and AspM17 via a single water molecule. However, we find that these Asp, which are both ionized in the MD trajectory, are not connected to each other even via four bridging waters. Rather AspL213 and AspM17 are key residues for proton transfer along two separate paths.

### The effects of mutating key nodes in the hydrogen bond network near Q_B_

The kinetics and thermodynamics of the coupled electron and proton transfer to Q_B_ have been well studied by mutation of residues near Q_B_ (Okamura et al. 2000a; Wraight 2004; Gunner et al. 2008). Proton uptake is not rate limiting at physiological pH in wild-type RCs so only changes that slow the proton transfer rate to be comparable with that of electron transfer will be visible (Eqn 1). A general finding is that mutation of residues closest to Q_B_ including SerL223 and AspL213 slow the proton uptake of the first proton, while mutation of GluL212 affects the delivery of the second proton. In contrast, mutations of individual residues in the middle of the network, including AspL210, AspM17, or GluH173 (Okamura et al. 2000a; Ishikita and Knapp 2005) or at the proton entry such as HisH126 or HisH128 do not strongly influence proton transfer. However, appropriately chosen double mutations have measurable effects on proton uptake.

### The role of SerL223

Every path in our network leads to Q_B_ via SerL223 (Ishikita and Knapp 2004b). Mutation of SerL223 to Ala or Asn slows proton uptake leading to a decrease in the rate of the second reduction of Q_B_ (Okamura et al. 2000a). In our network SerL223 plays an essential role as connecting all three proton transfer pathways to Q_B_. Mutations of SerL223 should require changes in proton delivery. Substitution of the Ser by Gly has a small effect on the reaction, which may be because the Gly leaves space for a water to reconnect the quinone to the network (Paddock et al. 1990).

### The role of GluL212

GluL212 is a key residue for delivery of the second proton to Q_B_. It is likely to be anionic in the Q_B_ ground state and it binds a proton in the presence of Q_B_^•-^. The proton binding has a pK_a_ of 9.8 so at higher pHs the second electron transfer to Q_B_ is slowed (Kleinfeld et al. 1984; Takahashi and Wraight 1992). Mutation of GluL212 to Gln does not affect the rate of the first reduction of Q_B_ but slows the second reduction by a modest amount (Okamura et al. 2000a). In the trajectory used here Q_B_ is neutral and GluL212 is ionized as appropriate in the ground state. The GluL212 connected to SerL223 via three waters in this network.

### The role of AspL213

The buried AspL213 is connected to paths starting at all three entries. In addition, its charge helps to tune the protonation states of the other residues near Q_B_ and so affects the electron affinity and thus the equilibrium constant for electron transfer from Q_A_^-•^ (Paddock et al. 1994b, 1998). Mutation of Asp213 to Asn slows the second electron transfer to Q_B_ sufficiently that proton uptake is now rate limiting (Rongey et al. 1993).

### The coupling between AspM17 and AspL210

In the network found here AspM17 and AspL210 are not connected to each other. M17 is on the path from Entry 1 and L210 is connected to Entry 2 and 3 (Figure 3). The fact that they are both ionized in the MD trajectory may lead them to move away from each other. However, support for their being on separate paths comes from the finding that single mutations of either residue has limited effects on the cycle of Q_B_ reduction (Okamura et al. 2000b). However, if both are mutated the first proton uptake becomes the rate limiting step, and there is also a decrease in the rate for uptake of the second proton (Paddock et al. 2001).

There is a well-studied cluster of residues on the surface including AspH124, HisH126, and HisH128. In the network presented here AspH124 is not a surface residue but it connects to HisH126, at E1, and HisH128 at E2. Mutation of any of these residues individually does not have a significant effect on the cycle of Q_B_ reduction, but mutation of both HisH126, and HisH128 leads to proton transfer becoming the rate limiting step, slowing the first electron and proton coupled second electron transfer (Adelroth et al. 2001). Crystal structures with Cd^2+^ or Zn^2+^ added show that these residues chelate the metal (Paddock et al. 2000; Adelroth et al. 2001). With metal bound to the wild-type protein proton transfer now becomes rate determining.

### Comparison of Q_A_ and Q_B_

While Q_B_ has many paths from the surface, Q_A_ is well insulated from the cytoplasm. The L polypeptide that surrounds the active branch bacteriopheophytin and then Q_B_ is 34% identical and 51% similar to the M polypeptide that surrounds the inactive branch bacteriopheophytin and Q_A_ (Williams et al. 1986). Each quinone is connected to the non-heme iron via a His. A count of the types of residues in the 10Å around each quinone finds they are not very different (Table SI.4). There are 13 charged and polar residues around Q_B_ and 14 around Q_A_. There are 25 hydrophobic residues in this volume around Q_B_ and 23 around Q_A_. However, counting the number of waters within 10Å of each quinone in the frames in the MD trajectory, finds there are 95.4±9 waters around Q_B_ and only 43.3±5 waters around Q_A_. In each frame there are 2.2±0.3 times more waters around Q_B_ than Q_A_.

Thus, similar residues are arranged to make quite different networks around the two quinones. The L and M were aligned (Figure SI.5). SerL223, which is the key connection from the network to Q_B_, is replaced with AsnM243 near Q_A_. Asn retains the polar character of the Ser, but the amide side chain cannot participate in Grotthuss proton transfer. AspL210 changes to Ala, HisL211 to Glu, GluL212 to Arg, AspL213 to Gly, Thr at L214 is conserved, while ThrL226 changes to Met. AspM17, which is an important residue on the path from Entry 1 to Q_B_ is in a region of the L polypeptide where BLAST reports no significant similarity to the M polypeptide.

Figure 5 compares the nature of the residues around the two quinones. Both quinones have surrounding hydrophobic residues (Figure 5A and 5C) as well as the charged and polar residues (Figure 5B and 5D). However, Q_A_ is complete surrounded by hydrophobic residues that forbid water entry (Figure 5A and 5B). In contrast, there is a well-defined slit that leads from Q_B_ to the surface along the polar and charged residues (Figure 5A and 5B). The polar residues directly around Q_A_ are needed to stabilize the anionic semiquinone, but they form an unconnected island. In contrast, the complex web of polar and charged region near Q_B_ attract waters into the network and are connected to the surface to facilitate proton uptake.

**Figure 5.**
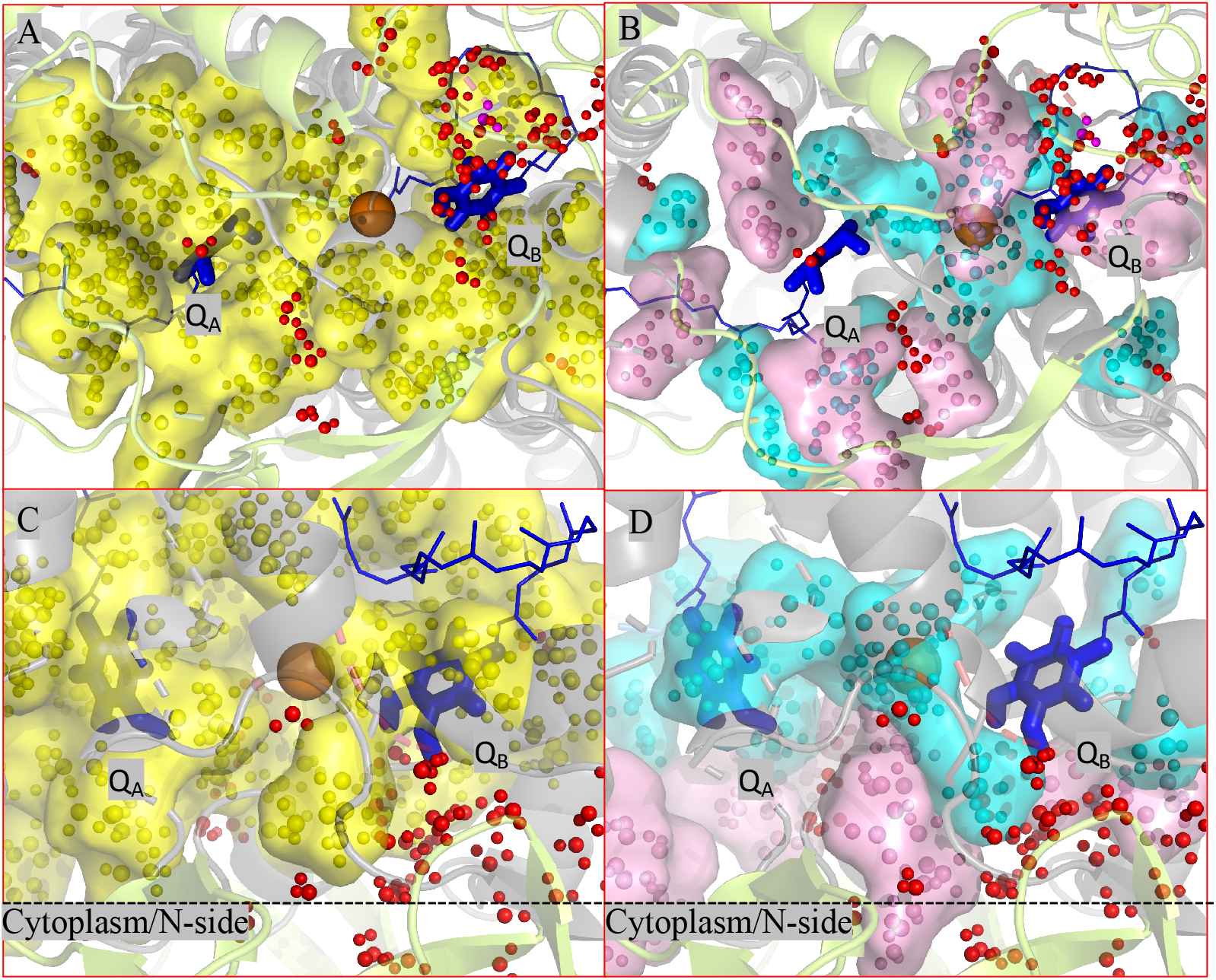
Surface showing hydrophobic and polar residues within 10 Å of Q_A_ and Q_B_. A and B look into the protein from the cytoplasmic surface. C and D are a side view with the cytoplasmic H subunit on the bottom of the figure and the dashed line showing the approximate position of the membrane. Q_A_ and Q_B_ are in blue, with the edge of the head groups visible. Water molecules are red balls. A and C: Hydrophobic side chains within the yellow surface. B and D: Charged side chains are in the pink surface and polar residues within the cyan surface.

### Limitations of the calculations

The networks represent an unbiased search for hydrogen bonded paths from the surface to Q_A_ and Q_B_. The residues of interest are not pre-chosen. The networks are permissive, with connections being drawn between pairs of residues in each MD frame, without consideration of their orientation or of the long-range order needed for Grotthuss proton transfers. As MD fixes the protonation states of all residues no changes in protonation states of intervening acidic and basic side chain are allowed. For example, some residues are suggested to be present in a distribution of protonation states in the MCCE calculations, such as GluL212, and GluH173, HisH126, HisH128, HisM301, LysH197, but they are ionized in the most populated state used to seed the MD trajectory. We expect that changes in residue protonation state and the addition of Q_B_^•-^ will lead to changes in the hydrogen bond network (Zhang et al. 2021). Future simulations will be carried out to answer the question how the network reorganizes to modify the likely proton path when the charge of the residues change and when the semiquinone is introduced.

The Q_B_ has long been recognized as having many possible pathways for proton transport from the surface (Sebban et al. 1995; Lancaster et al. 1996; Krammer et al. 2009b) as we find here. When there are many competing paths, there are additional considerations beyond the scope of this paper can help determine which could be preferred. These include the combining the frequency at which individual connections are made and the long-range connectivity of a path viewed from start to end. In addition, comparison of the free energy of a positive charge along different paths can show which favor the proton passage (Zhang et al. 2021; Kaur et al. 2021).

## 5. Conclusion

We have presented the hydrogen bond network that can transfer protons from the cytoplasm to Q_B_ while not to Q_A_. Three proton transfer pathways are found leading from three surface residue clusters to Q_B_ via residues and waters. In contrast, Q_A_ is well insulated from the protein surface. Our finding, in combination with earlier studies of the effects of mutation, shows the proton can likely traverse multiple paths to Q_B_. Thus, single mutations have much smaller effects than double mutations which change residues along different paths. For example, mutation of AspM17 or AspL210, which are located on different paths does not slow proton uptake, while mutation of both makes proton transport the rate limiting step. This shows the functionality of complex proton transfer networks in that they provides alternative paths and that longer and short paths can both be important.

## Supporting information

supplemental data Wei2022

## Acknowledgements

We acknowledge financial support from the Division of Chemical Sciences, Geosciences, and Biosciences, Office of Basic Energy Sciences, U.S. Department of Energy, Photosynthetic Systems. Computational studies grant DESC0001423. We thank the New York State’s Graduate Research Training Initiative (GRTI) for support of a high-performance computing cluster at the City college of New York used in this research. We thank Muhamed Amin and Gehan Ranepura for useful discussion.

