## supplemental data Wei2022 for "Comparison of Proton Transfer Paths to the Q_A_ and Q_B_ Sites of the *Rb. sphaeroides* Photosynthetic Reaction Centers"

**Part I:**

Figure SI.1 A: Structure used in Molecular Dynamics. B: RMSD backbone alignment within 150 ns MD trajectories.

**Part II:**

S.I.2, Distance restraints between the carbonyl oxygen of the quinone and the ND1 of HisH190.

**Part III:**

Table SI.3: Hydrogen bond connections with a maximum of 2 bridging water molecules in the network leading to Q_B_ in different time blocks of the MD trajectory.

**Part IV:**

Table SI.4: Residues found within t 10Å of the carbonyl oxygens of Q_A_ and Q_B_.

**Part V:**

Figure S.I.5: Blast comparison of L and M chains.

**Part VI:**

Links to the parameter and programs used here.

**Part I**

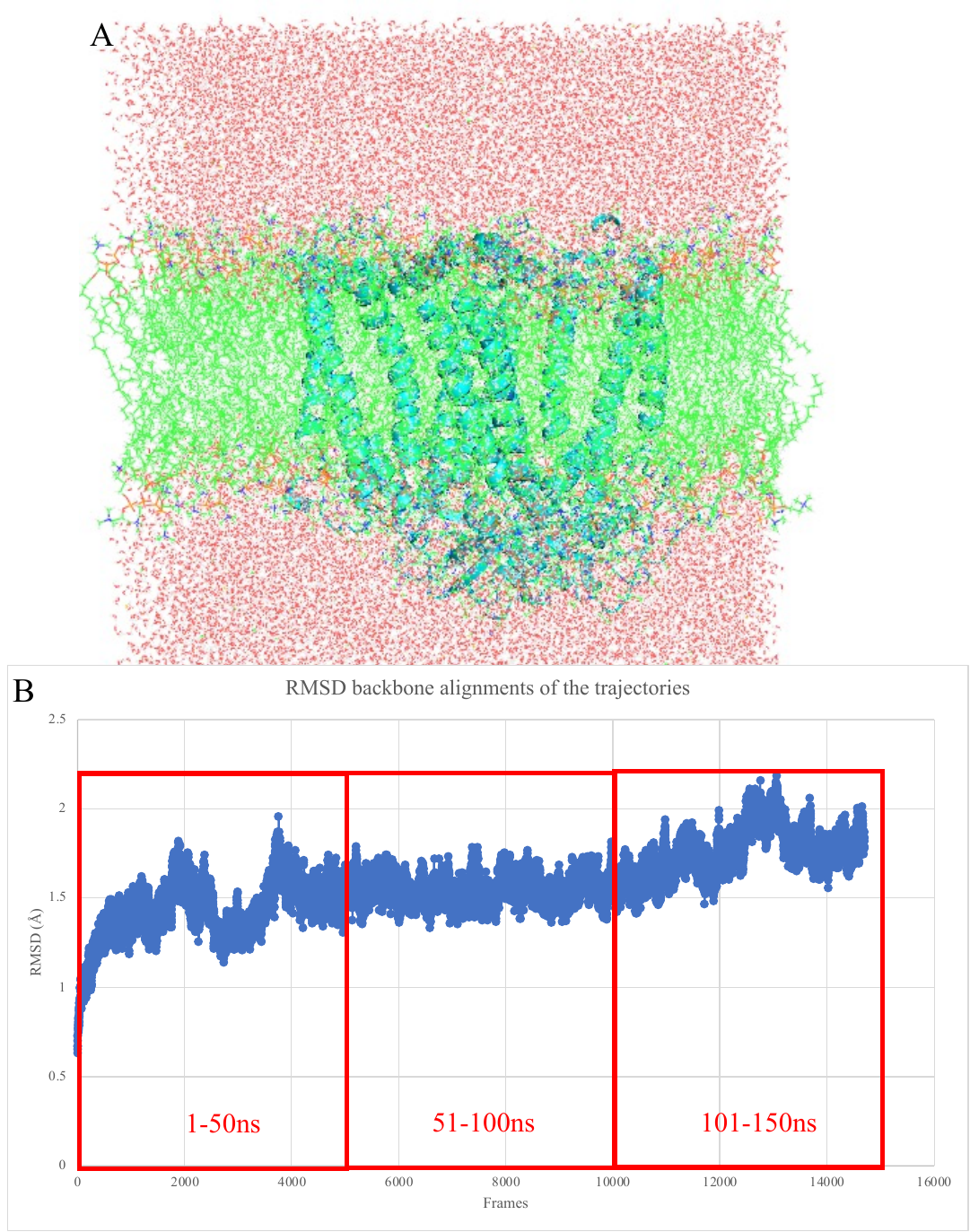

Figure SI.1 A: Coordinates used to start the MD trajectory are generated in the CHARMM-GUI. POPC lipids in green lines. Water molecules as red dots. RCSB (PDB ID: 1AIG provided the initial coordinates of the *Rb. sphaeroides* RCs. B. The RMSD backbone alignments of the trajectory. The network was derived from the 51-100 ns time block. The residues connectivity is maintained in the earlier and later portions of the trajectory. The average RMSD obtained from 150ns using VMD trajectory tool is 1.572±0.188 Å.

**Part II**

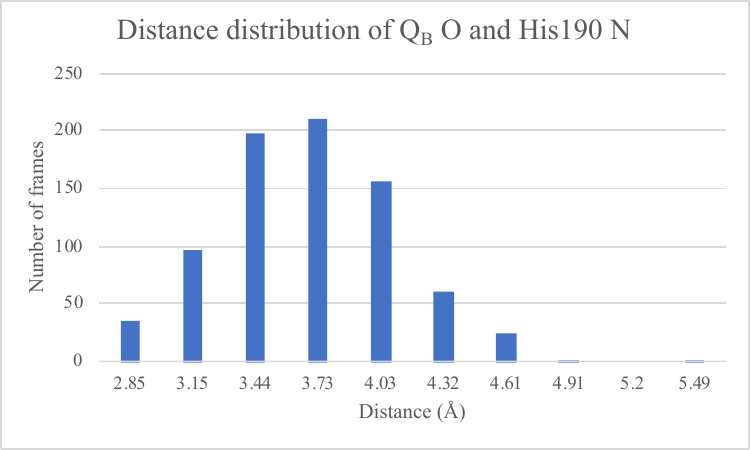

S.I.2, Histogram of the distance between the carbonyl oxygen of the quinone and the ND1 of HisH190. The unrestrained, neutral Q_B­_ will move from the proximal Q_B_ binding site toward the proximal site in the trajectory (Stowell et al. 1997). A 1000 Kcal/mol/Å constraint is placed between the O2(O_60 in MD) of Q_B_ and the ND1 of HisH190, which bridges the quinone and the iron. The average distance is 3.83±0.4. The inserted figure shows the variation in the distance with time.

**Table SI.3:** Hydrogen bond connections with a maximum of 2 bridging water molecules in the network leading to Q_B_ in different time blocks of the MD trajectory.

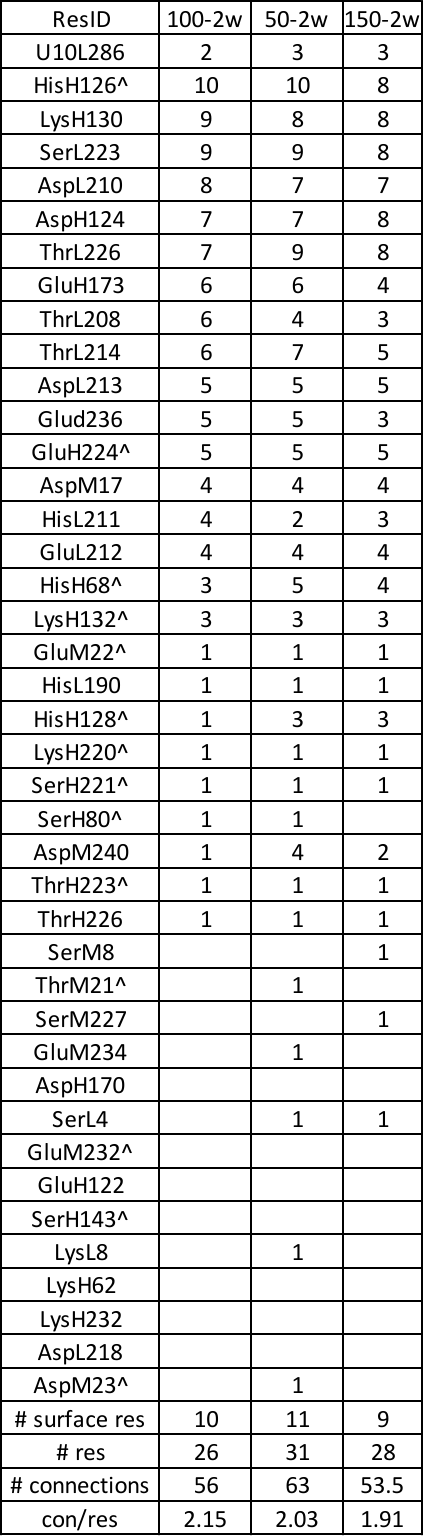

**Table SI.4**: Residues found within 10Å of the carbonyl oxygens of Q_A_ and Q_B_

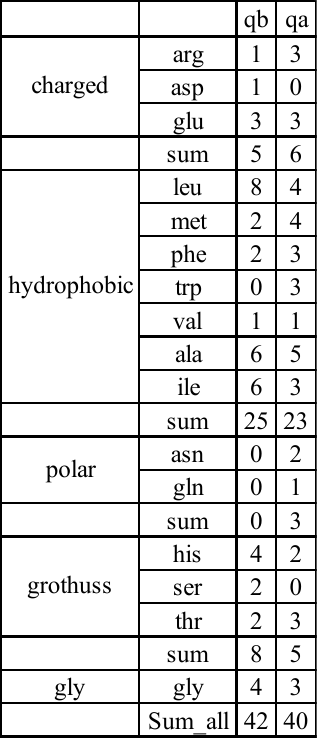

**Part V:** **Blast comparison of L and M chains**

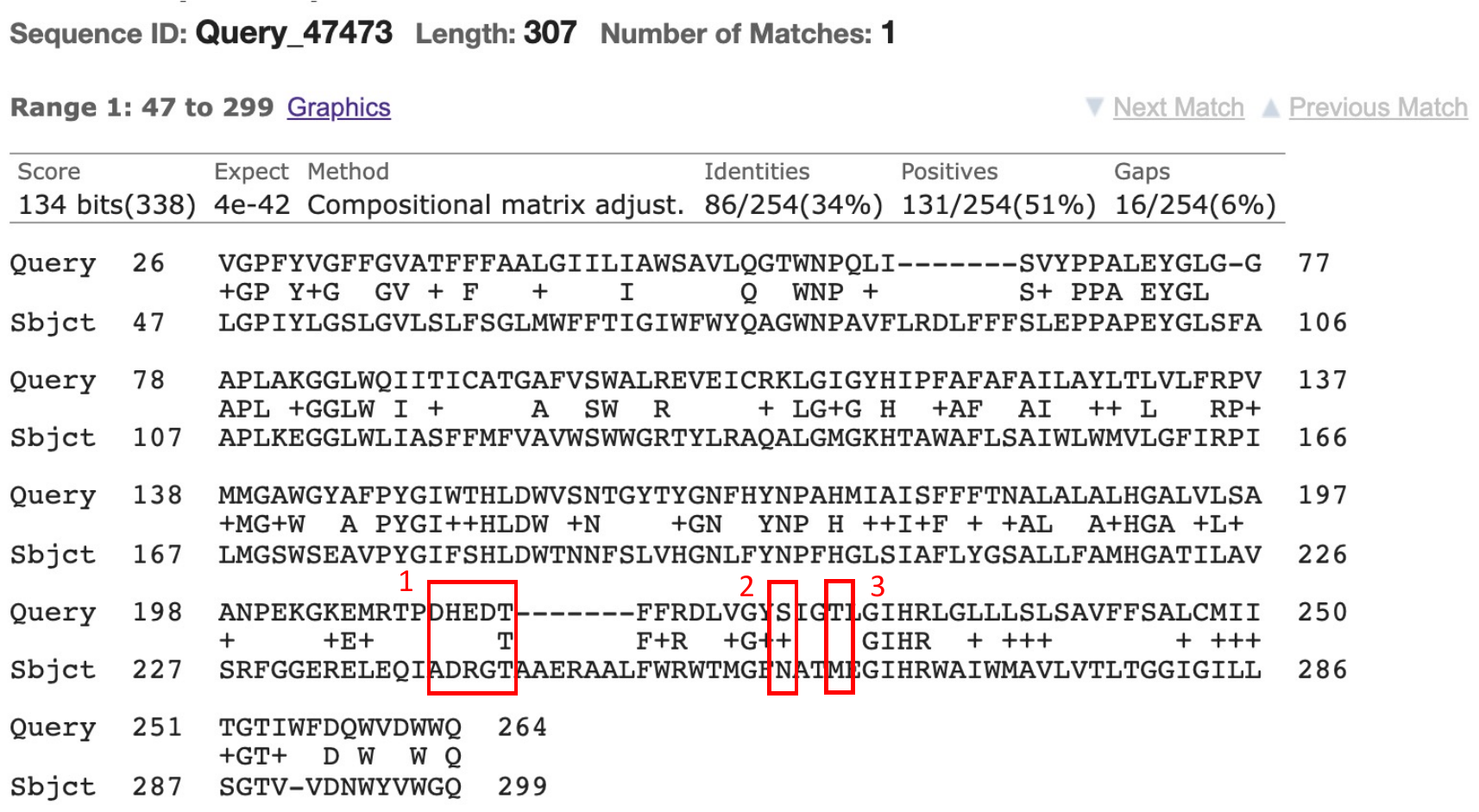

Figure S.I.4, Blast comparison of L (top) and M (bottom) chains. The residues in the network in the L chain and M chain that connect the surface to Q_B_ are marked in the red boxes. Box 1 includes residues from left to right: AspL210, HisL211, GluL212, AspL213, ThrL214. Box 2 is SerL223, and Box3 is residue ThrL226.

**Part VI: Programs used here can be found on GitHub**

1. The MCCE program is available at: <https://github.com/GunnerLab/Stable-MCCE>
2. The parameters taken from (Ceccarelli et al. 2003; LeBard et al. 2008), for the RC cofactors are at: <https://github.com/GunnerLab/RCs_parameters>
3. The script for stripping external water molecules while leaving those within the protein in an MD frame with explicit waters is at: <https://github.com/GunnerLab/Stable-MCCE/blob/master/bin/striph2o.py>
